# Long-range PCR amplification and nanopore sequencing of 8-10 kb mitochondrial fragments from environmental DNA

**DOI:** 10.64898/2026.01.13.699327

**Authors:** Stephanie A. Matthews, Olivia M. Scott, Yin Cheong Aden Ip, Elizabeth A. Allan, Ryan P. Kelly

## Abstract

1. Long-read sequencing data can provide increased taxonomic resolution and genomic linkage information that is otherwise impossible to obtain from short-read amplicon or shotgun sequencing. However, the use of long-read sequencing for environmental DNA (eDNA) analysis has thus far been limited by both the apparent rarity of long DNA molecules in eDNA samples and the lack of established bioinformatics workflows for long-read metabarcoding, particularly from mixed template samples.
2. Here, we report nanopore sequencing of 8.0 – 9.5 kb mitochondrial fragments obtained from mesocosm and field eDNA samples, amplified with long-range PCR (LR-PCR) using primers designed to preferentially amplify teleost mitogenomes.
3. We recovered half-mitochondria from 13 fish species in field-collected samples (Puget Sound in Washington State, USA), as well as from approximately half of the fish species inhabiting the positive control mesocosm (the Seattle Aquarium). Among biological replicates, we observed consistent detection of read-abundant species, while there was greater stochasticity in the presence of rarer species.
4. We demonstrate that long fragments can be obtained from standard eDNA samples and successfully amplified and sequenced to obtain species identifications despite higher error rates characteristic of nanopore sequencing. We present both laboratory methods and an accessible bioinformatic pipeline for obtaining and analyzing LR-PCR amplified fragments from eDNA, providing a framework for future long-read metabarcoding studies.

## 1. INTRODUCTION

Analyses of eDNA samples most commonly focus on amplification and sequencing of small fragment sizes due to the rapid breakdown of DNA in the environment and length restrictions in common sequencing platforms (Brandão-Dias et al. 2025). Previous attempts to isolate and sequence long fragments from field-collected eDNA samples have been limited to a few thousand base pairs (summarized in West and Deagle 2025). However, the DNA released by organisms includes intact mitochondria and larger fragments (Rodriguez-Ezpeleta et al. 2021; Mauvisseau et al. 2022) which can provide more information and might allow for greater species differentiation and haplotype detection (Doorenspleet et al. 2025). Because long fragments of DNA degrade quickly in the environment (Brandão-Dias et al. 2025), the detection of long fragments in a sample also provides a more recent snapshot of the community, lending spatiotemporal perspective to the shedding event.

Long-read sequencing of eDNA can be accomplished through direct sequencing of template DNA, either with or without size-selection and pre-processing steps (i.e., shotgun sequencing), or through targeted PCR amplification (i.e., metabarcoding) (Sigsgaard et al. 2020). Long fragments of DNA must be present in a sample for either method to accomplish long-read sequencing, but the methods differ in their sensitivity and scope of detection. PCR-based approaches allow for enrichment of target taxa or target fragment lengths, dependent on the primers used. In contrast, shotgun sequencing approaches provide a more comprehensive snapshot of the DNA present in the sample but can fail to detect long reads of interest due to rarer long metazoan fragments being overwhelmed by more abundant microbial sequences and more abundant small DNA fragments (Stat et al. 2017). Long-read sequencing platforms such as Oxford Nanopore Technologies (ONT) can capture kilo-base scale fragments in single reads, reducing dependence on bioinformatic assembly (Deiner et al. 2017), thus enabling detection of tandem repeats and other repetitive regions that complicate short-read assembly (Tørresen et al. 2019). For shotgun metagenomic applications, hybrid approaches combining long-read scaffolding with short-read polishing often achieve optimal results (Trigodet et al. 2022), but for targeted amplicon sequencing, long reads can provide complete gene or genomic regions in single molecules. Both PCR-enriched and PCR-free sequencing have been successfully used to recover full mitogenomes from single-species mesocosms (Mizuno et al. 2025), and PCR-enrichment has also enabled assembly of mitogenomes from short-read (300 bp) sequencing of field eDNA (Deiner et al. 2017).

Although standardized Illumina short-read metabarcoding workflows have become widely available (e.g., DADA2; APSCALE, Callahan et al. 2016; Buchner et al. 2022), analyzing ONT data remains challenging because raw error profiles and read-length heterogeneity complicate filtering, denoising, consensus, and taxonomic assignment (Delahaye and Nicolas 2021; Doorenspleet et al. 2025; Ip et al. 2025). Using ONT for short fragments is increasingly feasible, but community best practices are still diffuse. Long-read metabarcoding adds additional challenges (e.g., homopolymer indels, primer-derived chimeras, circular mitochondrial references, and length-bounded clustering). To harness useful information from long reads, bioinformatics must be explicit and reproducible.

As sequencing technology advances and it becomes tractable to routinely sequence longer fragments from environmental samples, it becomes more important to optimize methods for both eDNA laboratory workflows and bioinformatic analysis. Here we pair long-range PCR (LR-PCR) with ONT nanopore sequencing and a present a transparent pipeline to recover 8-10 kb fragments of fish mitochondria from field-collected eDNA samples (Puget Sound in Washington state), as well as from positive control mesocosm samples (Seattle Aquarium). Our workflow addresses the challenges of higher ONT error rates and high computational power required for long-read clustering by incorporating strict quality and length filtering criteria, utilizing consensus calling using amplicon_sorter (Vierstraete and Braeckman 2022), and BLAST-based taxonomic assignment. We demonstrate recovery and analysis of nearly half-mitochondrial fragments from eDNA samples and provide a practical methodological and analytical framework for long-read metabarcoding from mixed template.

## 2. MATERIALS & METHODS

### 2.1. Field eDNA collection and LR-PCR amplification

Field eDNA samples were collected on the April 2025 Salish cruise supported through the Western Ocean Acidification Center (WOAC). The samples were collected at 24 locations in the Salish Sea (Fig. 1, Appendix 1: Table S1). At each location, seawater was collected from approximately 10 m above the seafloor via CTD rosette. Once on board, triplicate 1-2 L samples were filtered directly from a single Niskin bottle onto a 5 μm MCE filter using an eDNA Citizen Scientist Sampler (Smith-Root, Vancouver WA) and preserved in 5 mL tubes each containing 1.5 mL of DNA/RNA Shield (Zymo). The preserved filters were stored at room temperature for approximately 11-16 days. DNA was extracted using a *Quick*-DNA Miniprep Plus Kit (Zymo Research), following all manufacturer’s protocols, and eluted in 100 μL of elution buffer. The concentration of total DNA in each sample was quantified using the Qubit 1X dsDNA HS Assay Kit with the Qubit 4 Fluorometer (Invitrogen) (Appendix 1: Table S1). The DNA extract was aliquoted into two 50 μL subsamples and one aliquot of each sample was stored at –80 °C for about 6 months, until used in this experiment.

**Figure 1.**
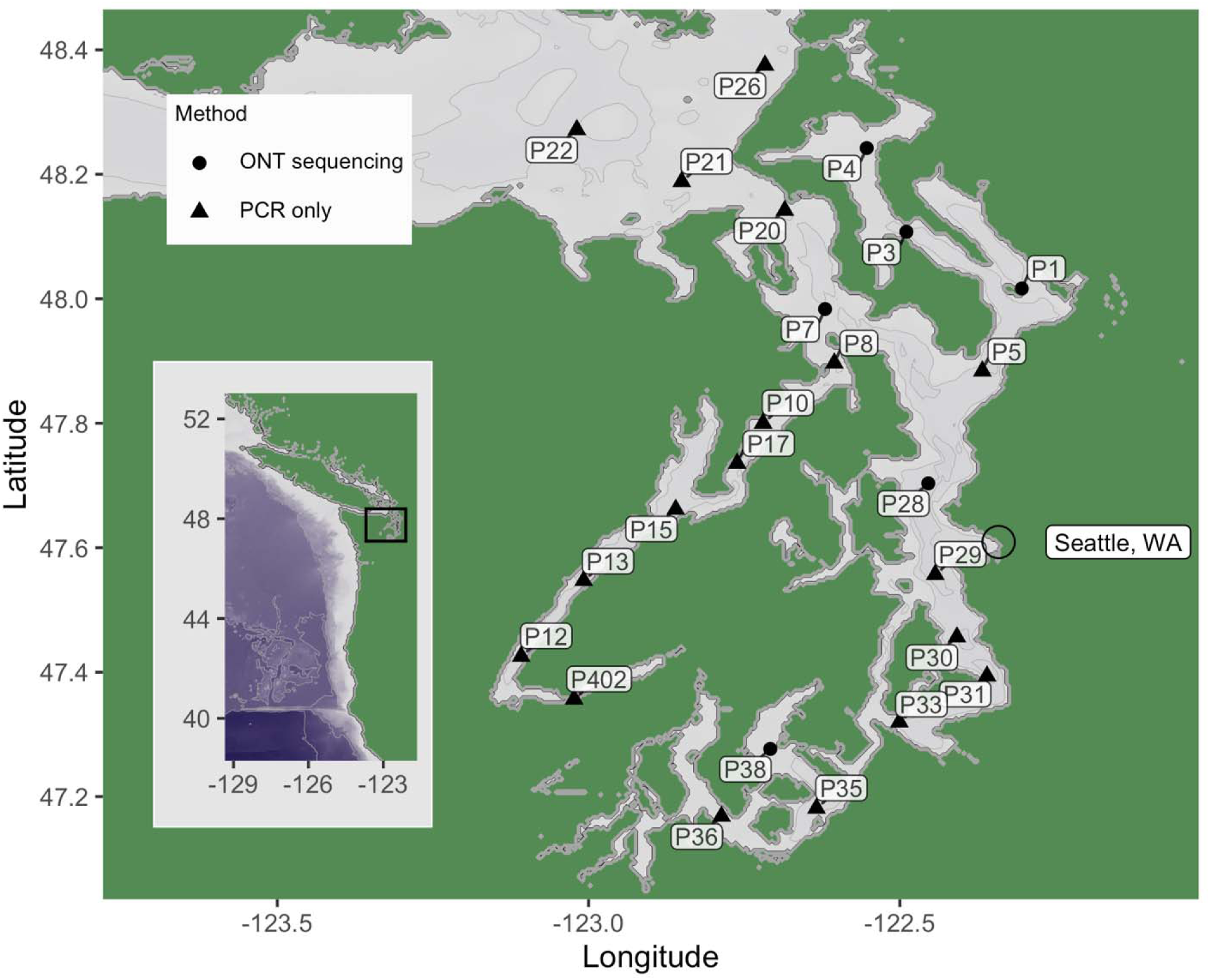
Geographic locations sampled in Puget Sound, Washington State, USA. Site names correspond to Salish cruise stations. Shape indicates whether the samples were sequenced or tested by gel imaging only. Inset shows the sampling map location relative to the west coast of North America. Seattle, WA is annotated for reference. Bathymetry and topography are sourced from the ETOPO 2022 global relief model from NOAA National Centers for Environmental Information. Bathymetry and topography data are freely available through the Creative Commons Zero 1.0 Universal Public Domain Dedication.

One set of triplicate mesocosm eDNA samples was included as a known-composition reference. The samples were collected from the Window on Washington Waters (WOWW) tank at the Seattle Aquarium in August 2024. The WOWW tank is approximately 454,000 L, contains 220 individual fish from 20 species (Appendix 1: Table S2), and has a wave machine that mixes the water. A clean 20 L Nalgene was submerged at the tank surface until filled, and three 1 L samples were each filtered onto a 5 μm MCE filter using an eDNA Citizen Scientist Sampler (Smith-Root, Vancouver WA). The filters were preserved in 5 mL tubes each containing 1.5 mL of DNA/RNA Shield (Zymo Research) and stored at room temperature until extraction. DNA was extracted within 24 hours of collection using the *Quick*-DNA Miniprep Plus Kit (Zymo Research), following all manufacturer’s protocols, and eluted in 100 μL of elution buffer. The concentration of total DNA in each sample was quantified using the Qubit 1X dsDNA HS Assay Kit with the Qubit 4 Fluorometer (Invitrogen) (Appendix 1: Table S1). The DNA extract was aliquoted into five 20 μL subsamples to minimize freeze-thaw cycles during use and stored at –80 ° C until used for this experiment, about 14 months after collection.

PCR amplification was performed using two previously published primers which preferentially amplify the teleost (ray-finned fishes) mitogenome (8.0 kb fragment: tRNA-Leu2(CUN) to 16S; 9.5 kb fragment: 16S to tRNA-Leu2(CUN)) (Table 1, Fig. 2) (Miya and Nishida 2000). These approximately 8.0 and 9.5 kb fragments constitute two halves of the mitochondria, excluding ∼27 bp in 16S. Each PCR reaction was comprised of 12.5 μL GoTaq Long PCR Master Mix (Promega Corporation), 0.625 μL of each 10 μM forward and reverse primer, 2 μL template DNA, and 9.25 μL nuclease-free water. To ensure careful handling of extracted eDNA, samples and PCR reactions were mixed by flicking or inverting, instead of being placed on a vortex mixer. PCR reactions were cycled for a 2 m initial denaturation at 94 °C, 27 cycles of 20 s denaturation at 94 °C and 10 m annealing/extension at 68 °C, and a final 10 m extension at 72 °C, following the manufacturer’s recommendation apart from modification of the annealing/extension temperature. A single technical replicate per biological replicate was visualized on a 0.7% agarose in TBE gel, with SYBR Green (Life Technologies Corporation) added to the loading dye for a final concentration of 1X, to determine presence or absence of target bands for all 72 field samples (Appendix 1: Table S1, Fig. S1). We note that the target bands visualized were offset from the ladder spacing, due to differences in loading concentration (Appendix 1: Fig. S2). As the sizes of the target bands were subsequently confirmed on the Genomic DNA ScreenTape Analysis (Agilent) (Fig. 3) and by the length of the ONT reads obtained, we did not attempt to re-visualize the bands using different techniques. Stations that had both 8.0 kb and 9.5 kb bands visible in all three samples were subsequently used for ONT sequencing (note: Station P38 was an exception with only 2/3 samples presenting the 9.5 kb band) (Table 2).

**Figure 2.**
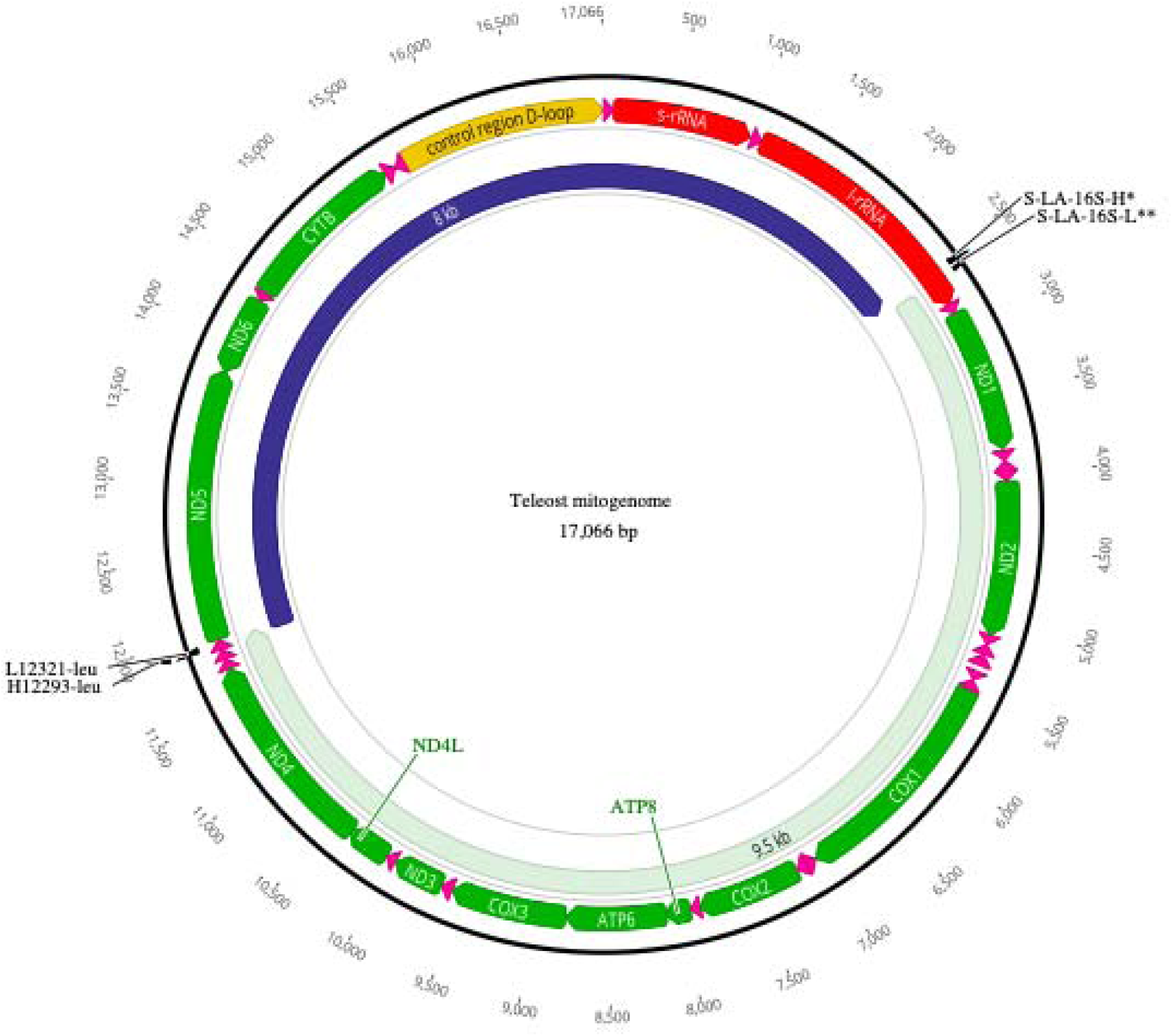
Map of a model teleost mitogenome showing the target fragments amplified by LR-PCR and nanopore sequenced. Amplicons are annotated with the expected length: dark blue shows the 8.0 kb fragment, light green shows the 9.5 kb fragment. Primer binding locations are indicated by primer names; primer sequences can be found in Table 1.

**Figure 3.**
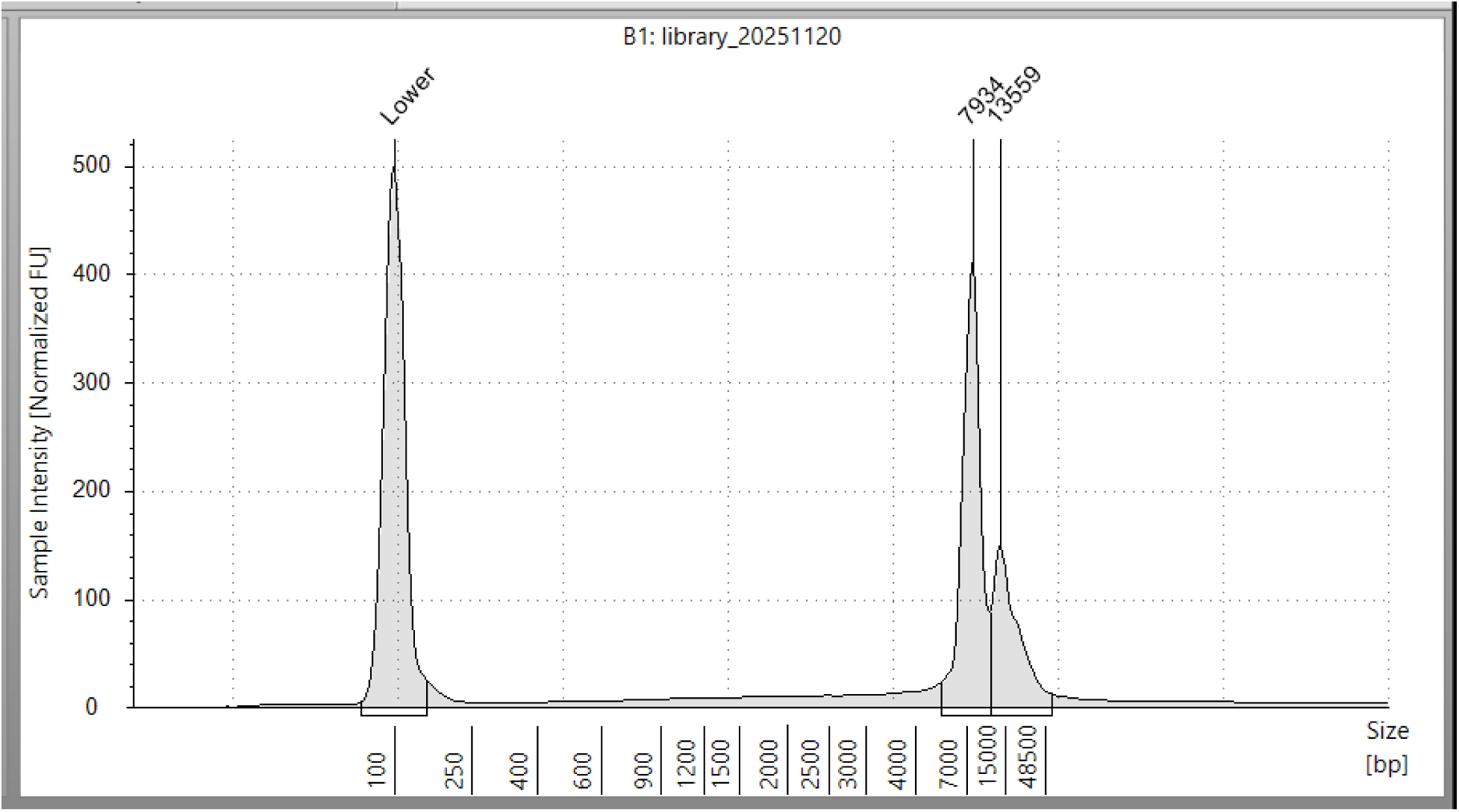
TapeStation gDNA ScreenTape Analysis electropherogram of the final ONT library showing the presence of 8.0 and 9.5 kb amplicon generated from eDNA samples from the aquarium and field samples.

**Table 1.**
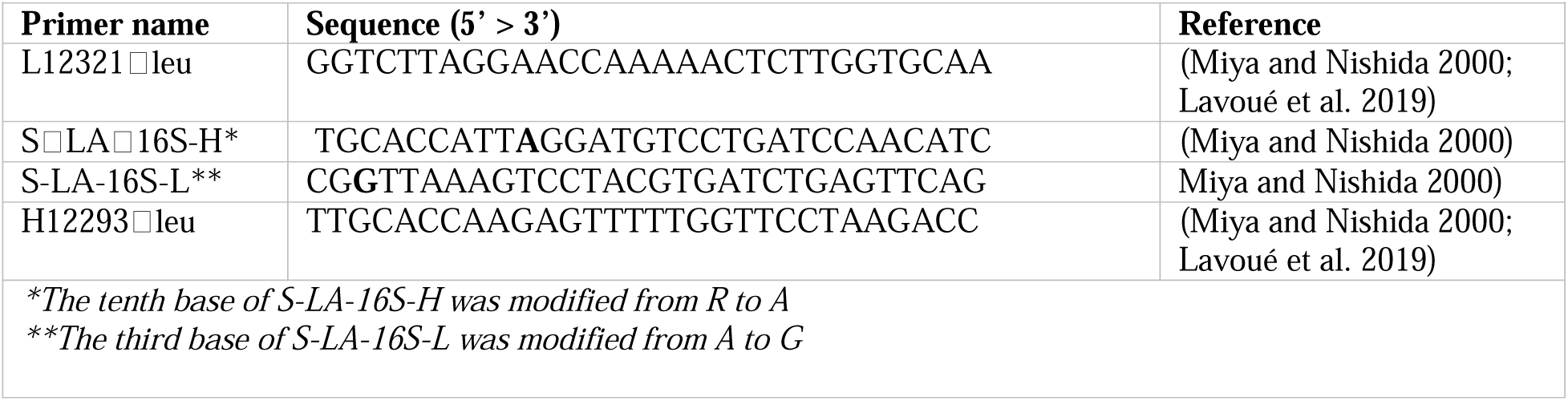
Long-range PCR primers used in this study.

**Table 2.** For each sampling location, the number of field sample replicates with visible target bands after LR-PCR. Field sampling locations subsequently used included in ONT sequencing are denoted by ^.

|  |  |  |
| --- | --- | --- |
| **Station** | **Number of samples with 9.5 kb band (out of 3)** | **Number of samples with 8.0 kb band (out of 3)** |
| P1 ^ | 3 | 3 |
| P3 ^ | 3 | 3 |
| P4 ^ | 3 | 3 |
| P5 | 2 | 3 |
| P7 ^ | 3 | 3 |
| P8 | 2 | 0 |
| P10 | 2 | 3 |
| P12 | 2 | 1 |
| P13 | 1 | 2 |
| P15 | 1 | 0 |
| P17 | 2 | 2 |
| P20 | 2 | 3 |
| P21 | 3 | 2 |
| P22 | 1 | 0 |
| P26 | 1 | 2 |
| P28 ^ | 3 | 3 |
| P29 | 0 | 1 |
| P30 | 3 | 1 |
| P31 | 0 | 0 |
| P33 | 1 | 1 |
| P35 | 0 | 2 |
| P36 | 1 | 2 |
| P38 ^ | 2 | 3 |
| P402 | 2 | 3 |

### 2.2 Library Preparation and Sequencing with ONT SQK-NBD114.24

To generate sufficient DNA to obtain 80-120 fmol of final library prepared with the Ligation Sequencing Amplicons - Native Barcoding Kit 24 V14 (SQK-NBD114.24) protocol (Oxford Nanopore Technologies), six replicate PCR reactions were run per target fragment for each eDNA sample, and three replicate PCR reactions were run per target fragment for each mesocosm sample. Replicates were pooled then bead cleaned with 0.7X AMPure XP Beads (Beckman Coulter) to remove DNA fragments < 250 bp in size. To increase yield during elution from the magnetic beads, samples were incubated at 37 °C for 60 min and pipette mixed with a wide-bore pipette tip every 30 min. To maximize the number of samples sequenced given the number of barcodes in hand, the 8.0 and 9.5 kb fragments were pooled together for each sample prior to sample ligation with a unique barcode, to be sorted subsequently (post-sequencing) by length and sequence identity. Pooled PCR products were quantified by using Qubit 1X dsDNA HS Assay (Thermo Fisher) and this value was used to estimate the moles of target DNA (assuming the average target size was 8750 bp) added to each barcoding reaction (Table 3), the entire amount of which was used for subsequent pooling. This approach essentially doubled our sample size, enabling us to sequence two target fragments from 24 samples using 24 commercially produced barcodes, rather than requiring 48 independent barcodes.

**Table 3.** Amplicons sequenced to obtain large fragments from eDNA. Total DNA indicates the Qubit value for each sample prior to LR-PCR. Demultiplexed ONT reads are reported for the two target fragments together, as fragments shared the same barcode within each biological sample. Unique amplicons were generated by amplicon_sorter and the target fragment identified based on mitochondrial annotation.

| Station | Biological Replicate | Volume Filtered (L) | Total DNA (ng/uL) | Approx. molecules of target added to barcode step (fmol) | ONT reads passing QC |
| --- | --- | --- | --- | --- | --- |
| P1 | A | 2 | 3.71 | 19.83 | 12,637 |
|  | B | 2 | 6.21 | 19.83 | 14,392 |
|  | C | 2 | 4.94 | 17.33 | 16,898 |
| P3 | A | 2 | 4.21 | 7.18 | 856 |
|  | B | 2 | 4.91 | 6.54 | 959 |
|  | C | 2 | 4.68 | 9.16 | 899 |
| P4 | A | 2 | 5.31 | 20.87 | 1,076 |
|  | B | 2 | 5.02 | 7.77 | 1,118 |
|  | C | 2 | 5.11 | 7.18 | 656 |
| P7 | A | 2 | 7.15 | 5.07 | 245 |
|  | B | 2 | 7.46 | 5.73 | 217 |
|  | C | 2 | 9.53 | 7.82 | 196 |
| P28 | A | 2 | 3.75 | 4.56 | 405 |
|  | B | 2 | 3.78 | 5.23 | 762 |
|  | C | 2 | 3.4 | 5.04 | 575 |
| P38 | A | 2 | 14.8 | 17.53 | 402 |
|  | B | 2 | 11.2 | 18.55 | 2,295 |
|  | C | 2 | 11.1 | 9.55 | 194 |
| Seattle Aquarium | A | 1 | 7.5 | 11.52 | 32,541 |
|  | B | 1 | 7.7 | 11.83 | 26,624 |
|  | C | 1 | 9.4 | 11.97 | 30,622 |
| NTC | A | 0 | *nd* | 0.00 | 0 |
| NTC | A | 0 | *nd* | 0.00 | 0 |
| EB | A | 0 | *nd* | 0.00 | 0 |

All parts of the protocol were followed as written, except for the bead clean-up steps after native barcode ligation and after adapter ligation where the ratio of AMPure XP Beads to sample was increased from 0.4X to 0.7X to increase DNA retention. The final library, consisting of the 8.0 and 9.5 kb barcoded fragments each from 18 field samples, three mesocosm samples, two no-template controls (NTC)s, and one extraction blank (EB), was validated for the presence of target bands using Genomic DNA ScreenTape Analysis (Agilent) (Fig. 3). The final Qubit reading showed that the mass of total DNA in the final library was 637 ng, or approximately 118 fmol of target DNA. The library was loaded onto a single MinION flow cell (FLO-MIN114) and sequenced for 17 hours until pore occupancy dropped below 5%. Raw nanopore sequencing data were acquired in POD5 format at hourly intervals, with reads filtered a minimum quality score of Q10. Live basecalling was performed using Dorado v0.9.1 using the Super-Accurate (SUP) model (5.0.0) on an MSI Raider 18 HX laptop equipped with an NVIDIA RTX 4090 GPU, managed via Windows standalone MinKNOW v25.05.12. Demultiplexing and barcode trimming were applied in real time during basecalling, and the resulting FASTQ files were exported for subsequent downstream processing.

### 2.4. Bioinformatic processing

Because of the relatively high sequencing error rate of ONT data relative to the more common Illumina data, we designed our bioinformatics pipeline to identify and retain high-confidence consensus sequences derived from the observed sequencing variants (Fig. 4). Basecalled reads were first filtered using seqkit (Shen et al. 2016) to retain only reads with mean quality scores greater than Q15 and with lengths between 7 and 10.5 kb. A minimum mean quality score of Q15 was selected to minimize any influence of sequencing errors on the consensus sequences generated. The length range 7 – 10.5 kb was selected to retain reads within approximately 1 kb of the expected length of the amplified fragments, to account for interspecific variability in mitogenome length and thus in fragment size.

**Figure 4.**
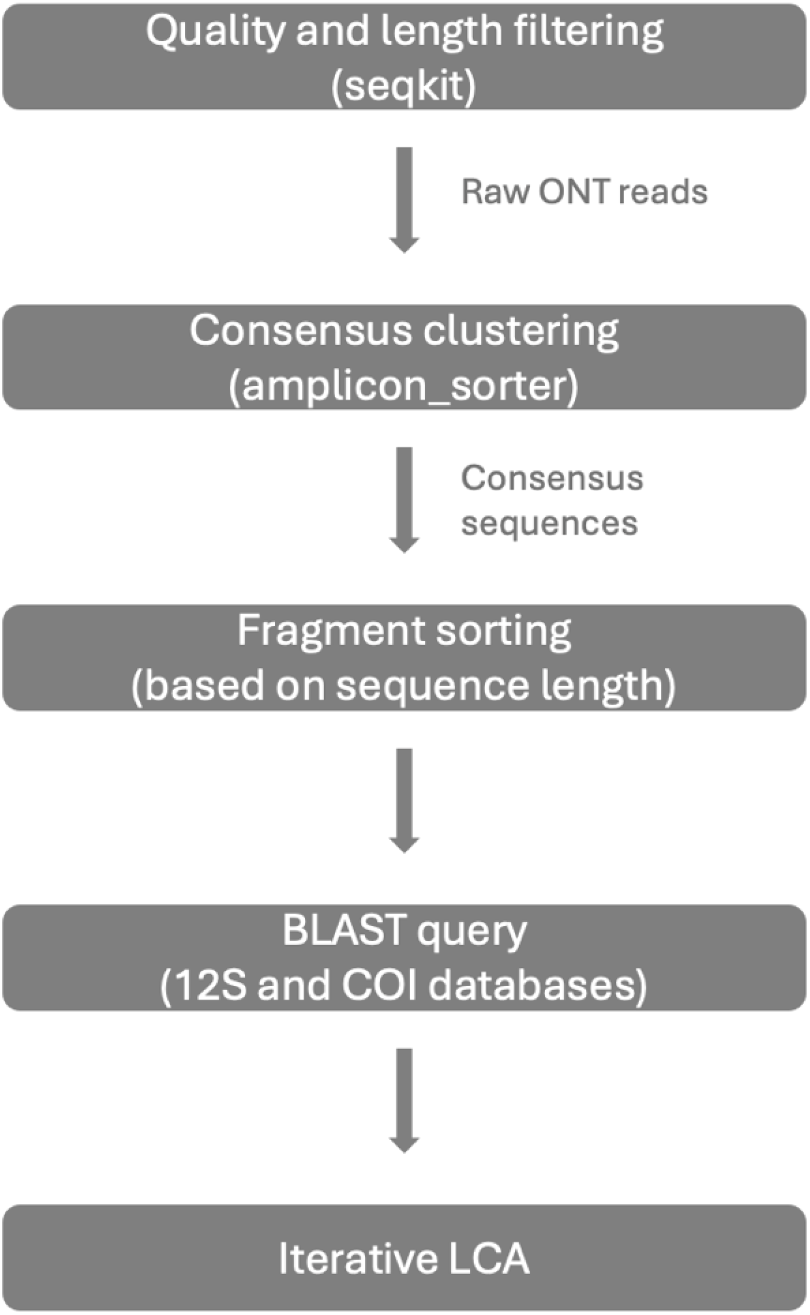
Flowchart showing the bioinformatic processing steps used to generate species identities from raw ONT reads. Detailed bioinformatic settings can be found in the text of the Methods section.

Filtered reads were clustered into amplicon groups using the program amplicon_sorter (Vierstraete and Braeckman 2022), a reference-free similarity-based clustering tool. The amplicon_sorter algorithm groups sequences by pairwise similarity without requiring reference sequences or a priori knowledge of expected amplicon composition, making it suitable for environmental samples with unknown species diversity. We used the settings all reads, comparison of all reads against each other, length_diff_consensus of 50, and all other defaults. Consensus sequences were treated as unique sequence variants, with the number of sequences contained within each group used as the read count for the sequence variant. Sequences not assigned to a group were excluded from subsequent analyses. We did not polish the final consensus sequences, as amplicon_sorter generates true consensus sequences, rather than identifying the centroid sequences of each cluster as is done by other ONT data processing algorithms (Vierstraete and Braeckman 2022).

To distinguish our barcoded half-mitochondrial fragments from one another, we employed a two-step bioinformatic process: first, barcoded reads were sorted into different fragment size bins; then, the reads within each bin were annotated with a taxonomic identity. For the first step, sequence variants were annotated for mitochondrial motifs in Geneious Prime using the annotated mitogenomes of *Oncorhynchus kisuch* (NC_009263), *Anarrhichthys ocellatus* (MG551528), *Clupea pallasii* (NC_009578), and *Hemilepidotus hemilepidotus* (NC_082813) as the reference database, with minimum 60% similarity required for annotation. Visual assessment of the annotated consensus sequences showed that the 9.5 kb fragment ranged from 9430 – 9520 bp, while the 8.0 kb fragment ranged from 7257 – 8951 kb. Based on these observed read lengths, sequences < 9.2 kb in length were classified as the 8.0 kb fragment containing 12S and sequences ≥ 9.2 kb were classified as the 9.5 kb fragment containing COI.

To assign taxonomy, amplicons were queried against custom 12S and COI BLAST databases generated using NCBI’s blastdb_alias tool on the core_nt database (downloaded 25 November 2025); such aliasing greatly decreases search times by only searching against a relevant subset of the larger database. The 12S aliased database was generated using GI numbers for database entries responding to search query “(Vertebrata[Organism]) AND (mitochondrion OR mitochondrial) AND (12s OR 12S OR \“complete genome\”) AND (0:20000[Sequence Length]).” The COI database was generated in the same way, using the search query “Vertebrata[Organism]) AND (mitochondrion OR mitochondrial) AND (COI OR \“cytochrome oxidase\” OR \“cytochrome c oxidase\” OR \“complete genome\”) AND (0:20000[Sequence Length]).” The GI numbers of all sequences included in these databases are provided as supplemental files for reproducibility (Appendix 2). Sequences were queried against the relevant database (12S or COI for the 8.0 and 9.5 kb fragments, respectively) using local command line megablast with the following parameters: -perc_identity 97 -word_size 100 -evalue 1e-10 - max_target_seqs 5. The BLAST hits were filtered to retain only results with ≥ 60% query coverage, to allow for matches that were split by the artificial linearization of mitogenomes in GenBank, with the control region / 12S boundary used as the linearization point. The lowest common ancestor (LCA) was calculated from the BLAST hits for each query sequence using taxonkit (Shen et al. 2016) in an iterative manner; if there were matches with 100% identity, LCA was calculated from only these, otherwise matches with ≥99.8, ≥99.6, ≥99.4, ≥99.2, ≥99.0, or ≥98 identity were considered, with cutoffs tested sequentially. The LCA was used as the final taxonomic assignment for the read, with two exceptions. Amplicons identified as *Hippoglossoides elassodon* (flathead sole), *H. robustus* (Bering flounder), or *Hippoglossoides sp*., due to BLAST hits with similar % identity to these two species, were all classified as *H. elassodon*, as *H. robustus* has been proposed to be a junior synonym of *H. elassodon* (Vinnikov et al. 2018). Similarly, amplicons identified only as *Hippoglossus sp.* due to BLAST hits to both *Hippoglossus stenolepis* (Pacific halibut) and *H. hippoglossus* (Atlantic halibut) were classified as *H. stenolepis*, as *H. hippoglossus* does not occur in the Pacific Ocean (Cargnelli et al. 1999). Read counts were summed within each species for each sample. As amplicon_sorter does not identify consensus sequences shared among samples, we used the LCA taxonomic assignment of each barcode to compare detections among replicates and across sampling locations. To account for potential barcode bleed among samples due to the NBD114 barcoding kit (Sauvage et al. 2023; Dai et al. 2025), we removed species from samples with within-sample occurrences < 3% of the total reads for the taxon. After quality, length, and taxonomic filtering, we obtained 46,010 long-read sequences from field eDNA samples (n=18) and an additional 83,492 long-read sequences from positive control eDNA samples (n=3). All code to reproduce this bioinformatic workflow is available at https://github.com/samatthews/LR-PCR_from_eDNA.

### 2.5. Statistical analysis

To calculate the average and standard deviation of reads obtained for field and aquarium samples, we used the summarize function from the R package *dplyr*. We also used this function to calculate the number of species observed in each station and within each LR-PCR fragment. We used a logistic regression to model the probability of detecting a species in the three biological replicates collected at each field sampling location, treating the two LR-PCR fragments independently (R package *stats*). All data visualizations were generated in R, with code to reproduce the analysis available at https://github.com/samatthews/LR-PCR_from_eDNA.

## 3. RESULTS

We queried 24 field sampling locations each with biological triplicates for a total of 72 samples for long fragments of eDNA using LR-PCR. At 23 of the locations (96%), we observed target-size bands in at least one of the six PCR reactions tested across the two size fragments (Table 2; Appendix 1 Fig. S1). Of the 72 unique field samples, 43 (60%) had positive detections of the 9.5 kb fragment and 46 (64%) had positive detections of the 8.0 kb fragment. Success rates of LR-PCR were higher for the aquarium positive controls than for the field samples: we observed target-size bands for both fragments from all three biological replicates collected and the bands observed on the agarose gel were brighter than for most of the field samples (Appendix 1 Fig. S1). We recovered more ONT long reads (passing quality and length filtering) from the samples with brighter bands, reflecting the fact that we did not normalize the amount of DNA per sample in the final library pool. We obtained on average 29,929 ± 3,019 reads per biological sample from the aquarium eDNA samples, and 3,043 ± 5,410 reads per biological sample from the field samples (Table 3).

From the 24 field samples screened for amplification, we selected six locations to sequence (each with three biological replicates). In total, in the six field sampling locations included in the ONT sequencing run, 13 unique species were observed, all of which were identified as Teleostei (Fig. 6). All observed species have biogeographic distributions that include Puget Sound and are expected to occur near the seafloor in muddy or sandy habitats. All 13 species were observed in the 8.0 kb fragment, and eight species were observed in the 9.5 kb fragment. In five of the six field sampling locations included for ONT sequencing, we observed 1-2 species from each fragment in each of the biological replicates (Fig. 5A). The exception was station P4, which had reads from 2-5 species in both fragments across all biological replicates. *Merluccius productus* (Pacific hake) was the species with the most reads observed in the LR-PCR sequences, appearing only at station P1. *Clupea pallasii* (Pacific herring) and *Gadus chalcogrammus* (Alaska pollock) were also relatively well-represented in the sequences, with *C. pallasii* appearing only at station P4 and *G. chalcogrammus* detected at P3, P4, and. P28. All other species were observed at lower read abundances. Six of these rarer species occurred only in a single sampling location, and four occurred at two locations.

**Figure 5.**
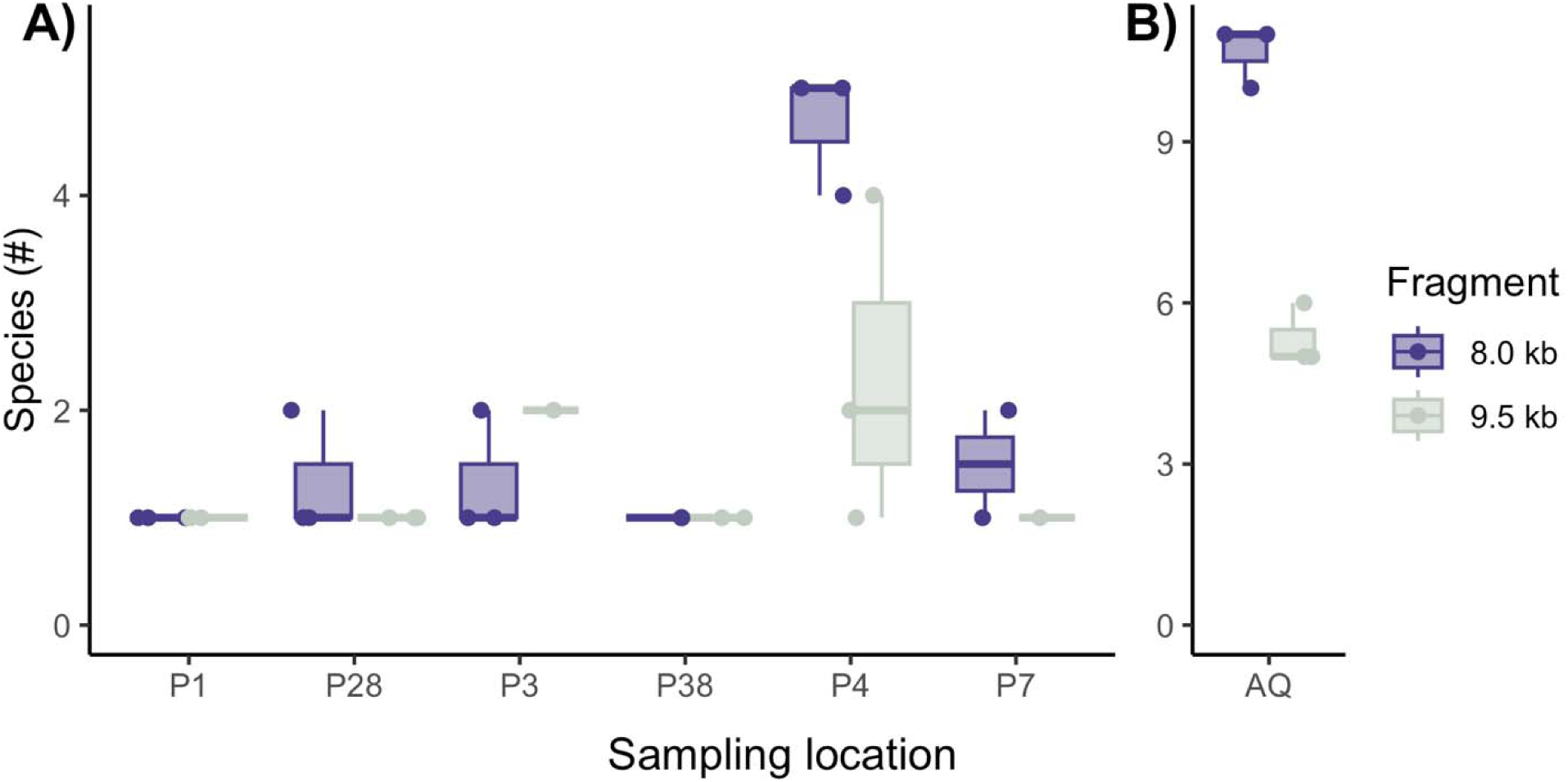
Richness (number of unique species) observed via LR-PCR and ONT barcoding at each sampling location. A) Field eDNA samples collected from Puget Sound, Washington state. B) Mesocosm eDNA samples collected from the Seattle Aquarium and sequenced as a positive control. Colors indicate the two target fragments amplified.

**Figure 6.**
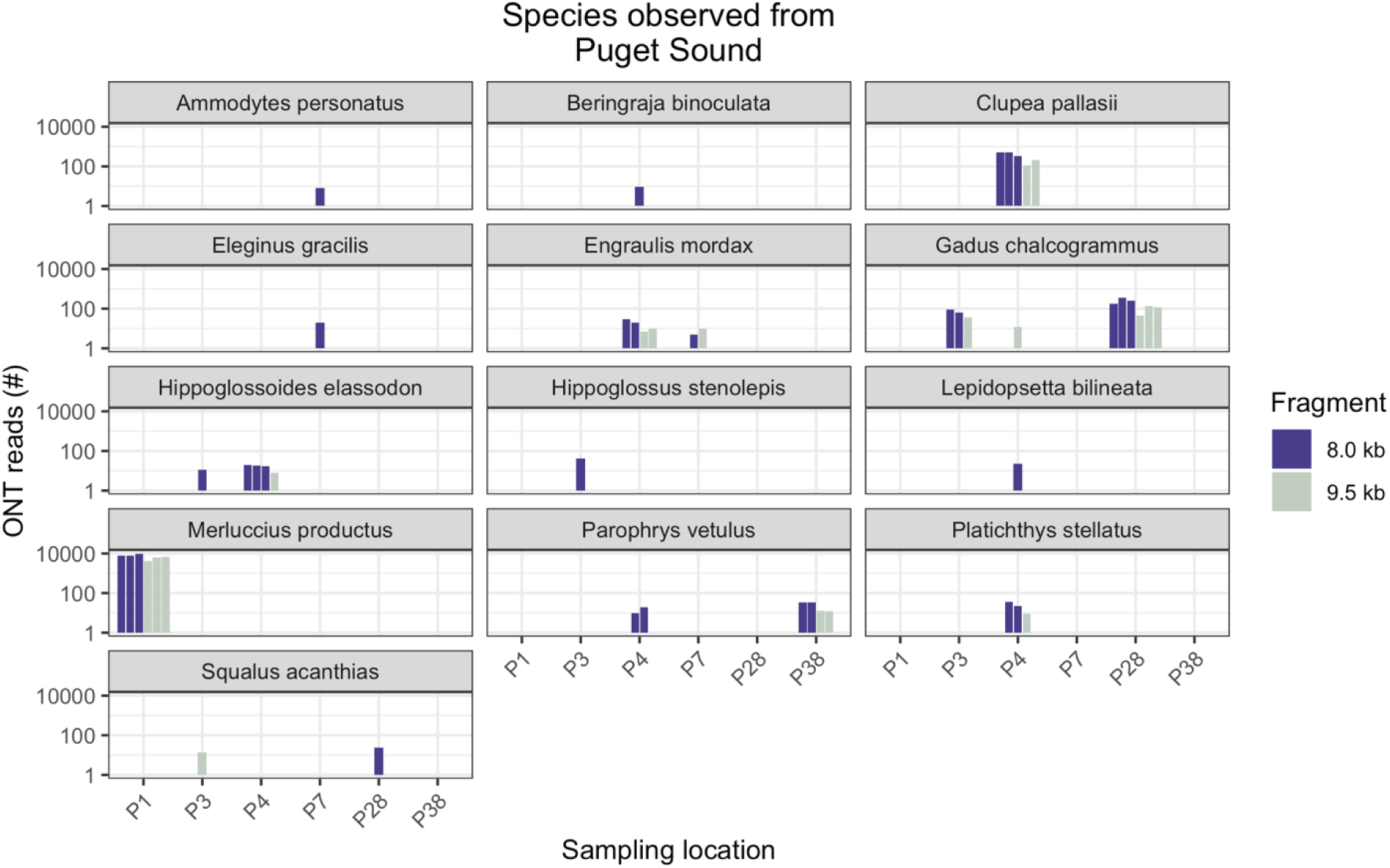
Species detected from Puget Sound using LR-PCR and ONT sequencing. Each panel shows one species observed in the sequence data, with the height of the bars (y-axis) indicating the number of reads assigned to the consensus sequence for each of three biological replicates at each sampling location (x-axis). The 8.0 kb fragment containing the 12S gene is shown in dark blue, the 9.0 kb fragment containing the COI gene is shown in light green.

The positive controls from the Seattle Aquarium had higher per-sample species richness than in field-collected eDNA samples, as might be expected from the very high fish concentration of the aquarium. There, we observed 10-11 species per biological replicate (13 total) in the 8.0 kb fragments, and 5-6 species per biological replicate (6 total) in the 9.5 kb fragment, for a total of 13 fish species in the two fragments (Figs. 5B, 7). As in the field-collected eDNA samples, all observed species were identified as Teleostei. We detected eight of the 20 fish species known to inhabit the mesocosm as well as five species not expected from the census of the aquarium tank (Appendix 1: Table S2). *Anarrhichthys ocellatus* (wolfeel), *Oncorhynchus kisutch* (Coho salmon), and *Sebastes flavidus* (yellowtail rockfish) had the most reads recovered from the aquarium sample, all of which were known to inhabit the tank. *S. melanops* and *S. pinniger*, both expected according to aquarium records, were also moderately abundant. The most abundant unexpected species was herring, *Clupea pallasii*, which is both common in the region and also in aquarium feed. Approximately 7-8% of the total reads recovered for each aquarium sample (including both fragments) were identified as *C. pallasii* (2,334 ± 296 reads)

**Figure 7.**
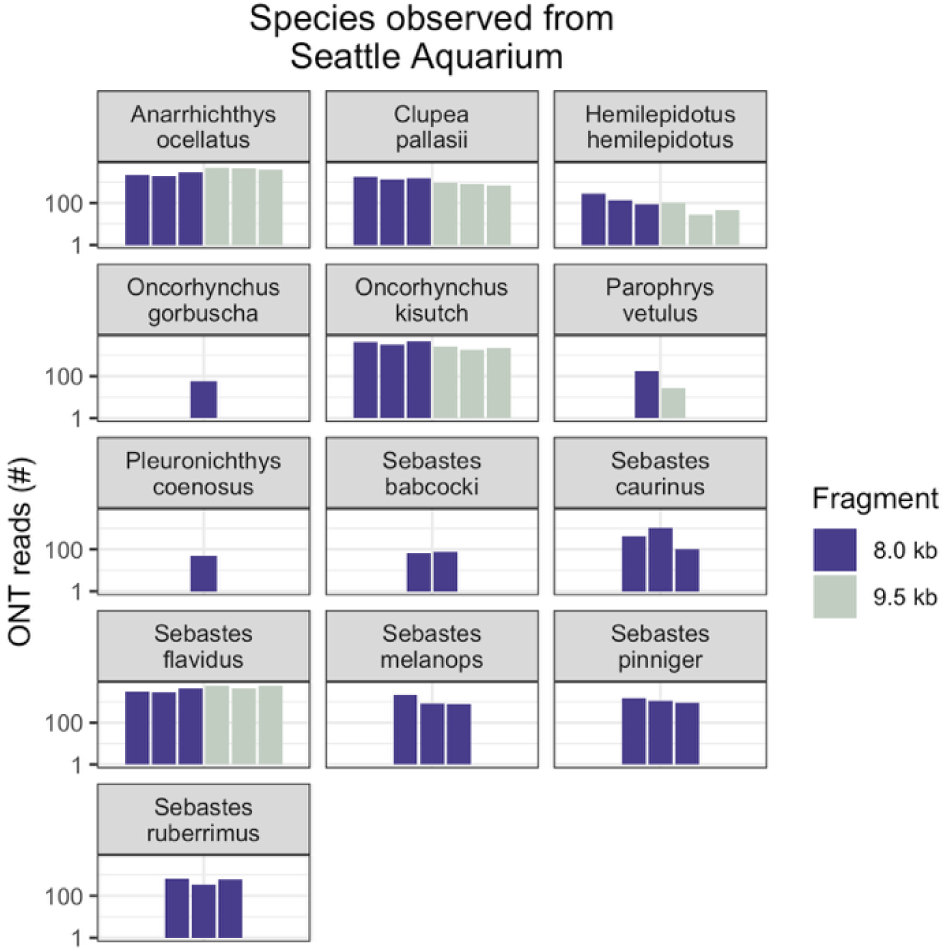
Species detected from the Seattle Aquarium positive control eDNA samples using LR-PCR and ONT sequencing. Each panel shows one species observed in the sequence data, with the height of the bars (y-axis) indicating the number of reads assigned to each species for each of three biological replicates at each sampling location (x-axis). Note log10 scaling of the y-axis. The 8.0 kb fragment containing the 12S gene is shown in dark blue, the 9.0 kb fragment containing the COI gene is shown in light green.

In both the field samples and the aquarium positive control, abundant species appeared in more biological replicates than rarer species (Fig. 8). Species with higher read counts (> 100 reads per sample) included *M. productus*, *C. pallasii*, and *G. chalcogrammus* in the field samples and *A. ocellatus, O. kisutch, and S. flavidus* in the positive controls. These species were typically observed in all three biological replicates and were detected by both the 8.0 and 9.5 kb fragments (Figs. 6, 7). In contrast, species with lower read counts (< 100 reads per sample) were often observed only in a single biological replicate, although some of these less abundant species were detected in multiple sampling locations (including *Engraulis mordax, Hippogossoides elassodon, Parophrys vetulus,* and *Squalus acanthias*).

**Figure 8.**
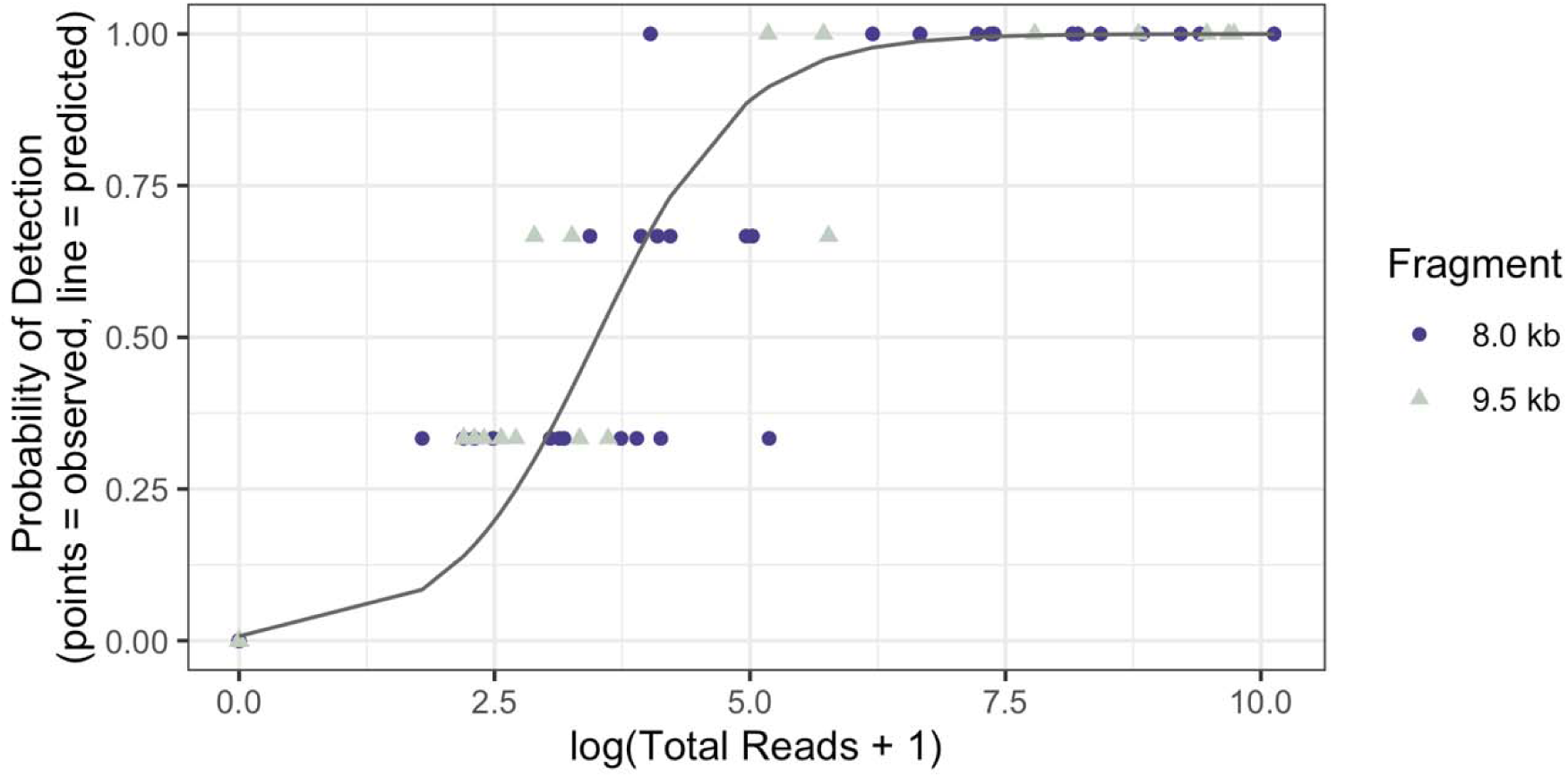
For each species and each LR-PCR fragment, the probability of detection in the three biological replicates collected at each field sampling location, as a function of the total number of reads obtained for the species across all three biological replicates. Data points are calculated independently for the two LR-PCR fragments; the 8.0 kb fragment containing the 12S gene is shown in dark blue and the 9.0 kb fragment containing the COI gene is shown in light green. The line indicates the predicted relationship, estimated from both fragments using a logistic regression.

## 4. DISCUSSION

We generated and sequenced amplicons of up to 9.5 kb from field-collected eDNA samples (Puget Sound in Washington state), as well as from positive controls collected from the Seattle Aquarium. Long-read sequencing on the Oxford Nanopore Technology (ONT) MinION platform detected 13 fish species from six field sampling locations, as well as 13 fish species from the aquarium. Our results confirm the presence of a multispecies reservoir of large eDNA fragments both in mesocosms and in field-collected samples, and provide a straightforward, repeatable protocol for using these long fragments for downstream eDNA analysis. This is the first demonstration of reproducible sequencing of 8-10 kb amplicons from eDNA field samples.

### 4.1. Methodological considerations for LR-PCR from eDNA

Compared to shotgun metagenomic strategies to recover long mitochondrial fragments (Deiner et al. 2017; Nousias et al. 2024; Mizuno et al. 2025), our LR-PCR method offers higher sensitivity for target taxa (via PCR enrichment) and simpler bioinformatics (no assembly required).

However, shotgun approaches provide genome-wide data and avoid PCR bias. The choice of method depends on the research question: LR-PCR is optimal for rapid, cost-effective assessment of a target community, while shotgun sequencing provides comprehensive biodiversity inventories including microbes and pathogens.

Although we did not exhaustively test all possible collection, preservation, extraction, and amplification steps that might impact the recovery of long reads, we note some strategies that may prove useful in future attempts to obtain long reads from eDNA. First, we used 5 μm filters for all of our sample collections, which may have enhanced the detection of large DNA fragments as larger pore sizes have been found to return greater proportions of large fragments (Jo et al. 2020). Because we expected rapid breakdown of large DNA fragments (Brandão-Dias et al. 2025), we preserved our sample immediately after filtration and extracted the DNA promptly to minimize microbial breakdown, which is reduced but not entirely inhibited by most common preservative methods (Cerk et al. 2025). Once extracted, we aliquoted DNA samples into multiple tubes and stored them at -80 C, to minimize freeze-thaw cycles which can fragment the stored DNA (Shao et al. 2012). Spin-column extraction and pipette shearing have previously been reported to negatively impact HMW DNA (Jaudou et al. 2022; Trigodet et al. 2022), but we found that spin-column based extraction methods were sufficient for recovery of our target amplicons and we did not use wide-bore pipettes for routine sample handling, although extracted eDNA samples and PCR reactions were flicked or inverted instead of vortexed.

### 4.2. Bioinformatic Considerations for Long-Read eDNA metabarcoding

The analysis of long-read amplicon data from environmental samples presents distinct challenges compared to established short-read workflows. The higher error rates of nanopore sequencing (6-10% vs <1% for Illumina; Delahaye and Nicolas 2021) require careful consideration of quality filtering thresholds, consensus calling approaches, and taxonomic assignment criteria, particularly with mixed template samples. In our pipeline, we addressed these challenges through several key steps: (i) stringent quality (Q>15) and length (within ∼10% of expected fragment length) filtering to remove the lowest-quality reads while retaining sufficient data for consensus building, (ii) reference-free similarity-based clustering with amplicon_sorter (Vierstraete and Braeckman 2022) to group similar sequences without requiring a priori taxonomic knowledge, and (iii) conservative taxonomic assignment requiring >98% similarity and high query coverage (>60%) for species-level identification.

For sequence clustering, we used amplicon_sorter (Vierstraete and Braeckman 2022) rather than statistical denoising algorithms designed for Illumina data (e.g., DADA2; Callahan et al. 2016). Statistical denoisers model sequencing errors based on platform-specific error profiles, and existing models are trained on Illumina data with <1% error rates. Given nanopore’s error profile (6-10% error rate with different error modes including varying flowcell chemistries, homopolymer miscalls and indels), we opted for consensus-calling using similarity-based clustering. This approach groups similar reads without assuming a specific error model, allowing the consensus sequence to emerge from multiple reads per amplicon. Future development of nanopore-specific denoising algorithms would be valuable, particularly for low-coverage samples where consensus calling from few reads may be unreliable.

To maximize the number of samples we could include in the 24-barcode kit, we did not use unique ONT indices to multiplex the two amplicons within each biological sample. Rather, we exploited the non-overlapping nature of the two amplified fragments to identify unique consensus sequences. Together with similarity-based annotation of mitochondrial gene regions and identification of the range of fragment lengths observed across different species, this allowed us to differentiate reads originating from the two amplicons. While non-standard, our approach allowed for recovery of reads from 50% less amplified PCR than would have been needed for independent barcoding of the two amplicons, as well as inclusion of twice as many samples using the ONT Native Barcoding Kit 24 V14. We suggest that similar strategies may be useful when working with limited template or difficult-to-obtain field samples. We used the BLAST-based annotation tool available in Geneious Prime, but this annotation could also be accomplished through freely available tools such as MITOS or MitoAnnotator (Bernt et al. 2013; Zhu et al. 2023). We also note that the length ranges we used (±1 kb from the expected fragment lengths) reflect the variability we saw across reference mitogenomes for an amplicon of 8-10 kb, and including the variable length control region. Other fragments (of different lengths or targeting a different region of the genome or mitogenome) will have assay-specific length ranges that should be selected based on expected variability.

A critical challenge we encountered was the computational handling of circular mitochondrial genomes, which are represented as linear sequences in reference databases with arbitrary start positions. Our 8.0 kb amplicon spanned the standard mitochondrial database indexing start position (the D-loop / 16S junction), resulting in artificially truncated and duplicated BLAST hits where the query sequence aligned to both the 5’ and 3’ ends of the reference sequence. We note that virtual PCR programs often fail due to the same reason, making it difficult to test LR-PCR primers *in silico*. This issue is likely to be encountered in any long-read mitochondrial metabarcoding study and should be considered in pipeline design, either through manual inspection or by pre-circularizing reference databases (e.g., Mitos2, Bernt et al. 2013; GetOrganelle, Jin et al. 2020).

### 4.3. Multiple vertebrate species contribute long eDNA fragments

While full mitochondria have been assembled from mesocosm eDNA, including recovery of full-length mitochondrial sequences (Mizuno et al. 2025), the presence of long DNA fragments in marine environments is not well documented, and the recovery of long reads from field-collected eDNA remains challenging. Our reported sequencing of long eDNA fragments from field-collected eDNA is a substantial advance, given the rapid decay of long DNA fragments (Brandão-Dias et al. 2025) and the expected rarity of long fragments in field-collected seawater (West and Deagle 2025). Long-read sequencing has been used successfully in non-marine environments both with PCR amplification (4,500 bp, Jamy et al. 2020) and with PCR-free shotgun sequencing (average ∼1,000 bp, Koda et al. 2023; lengths not reported, Nousias et al. 2024, 2025). In marine environments, PCR-free long-read sequencing returns primarily non-metazoan targets (Patin and Goodwin 2022). PCR amplification allows for enrichment of metazoan long reads from eDNA (Deiner et al. 2017; Mizuno et al. 2025; West and Deagle 2025), but long fragments must still be present at much higher concentrations than short fragments for reliable detection, due to decreased efficiency of LR-PCR (West and Deagle 2025) and due to the rarity of long eDNA fragments relative to short ones (Brandão-Dias et al. 2025).

We obtained LR-PCR amplicons from 13 fish species in the eDNA samples collected from the Seattle Aquarium mesocosm and from 13 fish species in the eDNA samples collected from Puget Sound. The maximum richness we observed was 5 species within a field sample replicate, and 11 species within an aquarium-collected sample replicate. The multiple species observed in each sample indicates that there were various >= 8.0 kb fragments of DNA present in the eDNA samples. Previous comparison of species richness across metabarcoding amplicons of different sizes has found lower richness for longer fragments (Doorenspleet et al. 2025).

Lower richness in long fragments likely reflects the shorter timeframe in which that DNA could have been released to the environment, given the rapid decay rates of long DNA fragments in the environment (Brandão-Dias et al. 2025). We did observe more species in the 8.0 kb amplicon than in the 9.5 kb amplicon, perhaps aligning with previous observations of greater diversity in smaller fragments (Doorenspleet et al. 2025), although given the small sample size in hand and narrow range of amplicon lengths, it is difficult to know whether this effect is a result of sampling variability.

There are multiple potential sources for the LR-PCR reads assigned to species that were not expected to occur in the aquarium sample. *Clupea pallasii*, the most abundant unexpected species, is commonly used as an aquarium feeder fish. We do not have any records of the timing of feedings relative to our water sampling event, but we expect that these reads originate from feeding events, due to their abundance. The four remaining unexpected species (*Oncorhynchus gorbuscha, Parophrys vetulus, Pleuronichtys coenosus,* and *Sebastes babcocki*) were represented by approximately 100 reads or less in a single biological replicate. All four species are either found in other tanks within the aquarium or can be found in Elliot Bay, Puget Sound where the aquarium inflow water is sourced (approximately 3,000 L per minute flows directly into the Window on Washington Waters tank).

### 4.4. Lower sensitivity of LR-PCR than of short-read amplification

While we do not have comparable short-read metabarcoding data from our eDNA samples, the expected inventory of the 454,000 L Seattle Aquarium tank gives some estimate of the sensitivity of our LR-PCR and ONT sequencing approach. The primers used were designed to amplify all teleosts (Miya and Nishida 2000; Lavoué et al. 2019). Considering only the fish species listed in the aquarium inventory, we observed 8 of the 20 available species while 12 were undetected by our LR-PCR assay. The absence of over half of the expected species from our dataset, together with intermittent detection of some species (e.g., *E. mordax, H. elassodon, P. vetulus,* and *S. acanthias*) suggest that the concentration of species-specific DNA template for many species was below the limit of detection. That is, absence of a species from a LR-PCR dataset does not confirm a true absence from the environment.

We do not currently have data in hand to determine the minimum template DNA concentration needed for successful LR-PCR. We obtained positive agarose gel bands from samples with total DNA concentrations ranging from 3.4 – 22.6 ng μl^-1^ (mean 7.52 ng μl^-1^), while environmental samples with concentrations ranging from 1.6 – 14.8 ng μl^-1^ (mean 5.6 ng μl^-1^) either failed to generate any bands or were successful only for one of the two fragments we attempted to amplify (Table S1). However, total DNA concentration does not provide the concentration of the target fragment that would be required to estimate the sensitivity and detection threshold of the assay.

Despite uncertainty regarding the absolute concentration of template required for successful LR-PCR, the patterns we observed among replicates align with our expectations for how eDNA behaves in environmental samples. The intermittent detection of species with lower read counts and the ubiquity of species with higher read counts is consistent with sampling theory, in which higher-concentration fragments are more likely to be sampled vs. lower-concentration fragments, both during the sampling stage of taking biological replicates from the environment (observed in the variability among biological replicates) and during the subsampling stage in which small volumes of extracted DNA are used as aliquots in individual PCR reactions (observed in the variability between fragments). There are also possible effects of primer bias, which would differentially affect the amplification efficiency of species. While the original primers were designed to amplify teleosts (Miya and Nishida 2000), we did not test whether their binding efficiency varies among teleost species. We also changed one base of each primer, removing any degenerate bases, which likely influenced which species are well-amplified by the primer set. We expect any primer bias would decrease the observed template availability and could result in lower read abundance and greater variability among replicates.

The non-detection of over half of the expected species from the aquarium tank aligns with our expectations that long DNA fragments are rare in eDNA (Brandão-Dias et al. 2025), and LR-PCR has decreased efficiency relative to short-read PCR (West and Deagle 2025). The combined effects of rarer template and lower assay efficiency limit the applicability of LR-PCR for oligotrophic environments as well as for rare species. Based on our results in hand, we recommend a minimum of three biological replicates and higher volume filtration (> 2L) (as performed here), as well as PCR replicates (which were not tagged independently in our study).

## 5. Future work

Our results demonstrate that nanopore sequencing can successfully generate species-level identifications from long amplicons from environmental eDNA samples. The single-molecule nature of nanopore reads preserves linkage information across the entire amplicon regions and resolves repetitive regions that are difficult or impossible to assemble, such as the mitochondrial D-loop. Immediate opportunities for advancing LR-PCR from eDNA include controlled mesocosm experiments with known species abundances to calibrate read counts to biomass relationships (quantitative validation), design of degenerate primers with in silico validation against broader taxonomic groups (elasmobranchs, agnathans), time-series sampling in a range of environmental conditions to assess persistence of long fragments vs. short fragments in marine environments, and demonstration of intraspecific variation detection (haplotype resolution).

After establishment of baseline success, quantitative assays (e.g. ddPCR) could be used across a range of fragment sizes to establish relationships between species abundance, DNA shedding rates, and LR-PCR detection probability for different taxa. These developments will transition LRPCR eDNA from a proof-of-concept to an operational tool for marine biodiversity monitoring.

## Supporting information

Appendix2 (zipped GIs)

## Data availability

Oxford Nanopore Technologies sequencing data will be made available as basecalled fastq files through Zenodo upon acceptance for publication.

**Supplementary Files** Appendix1_TablesFigures_LRPCR_20251222.docx Appendix2_Reference_GIs.zip

## Acknowledgements and Funding

The Seattle Aquarium provided access to mesocosm tanks used for mesocosm testing. Field sample collection was supported by the Washington Ocean Acidification Center. Computational resources for bioinformatic analysis were supported through the Center for Environmental Genomics at the University of Washington. We thank Zachary Child and Anna Boyer for facilitating field sampling. Hiroki Yamanaka provided valuable advice in selecting primers for this experiment. This work was funded by Oceankind. The funders had no role in study design, data collection and analysis, decision to publish, or preparation of the manuscript.

## Author Contribution Statement

SAM, OMS, YCAI, EAA, and RPK conceived the ideas and designed methodology; SAM and OMS collected the data; SAM, YCAI, and RPK analysed the data; SAM and OMS led the writing of the manuscript. All authors contributed critically to the drafts and gave final approval for publication.

