## Appendix1 (Tables & Figures) for "Long-range PCR amplification and nanopore sequencing of 8-10 kb mitochondrial fragments from environmental DNA"

*Appendix 1 for:*

**Long-range PCR and nanopore sequencing of 8-10 kb mitochondrial fragments from environmental DNA**

**SUPPLEMENTAL TABLES**

| **Table S1**. eDNA samples collected from the Puget Sound, Seattle, WA, USA. Filters were preserved immediately after collection and stored for approximately 2 weeks before DNA extraction. DNA concentrations were measured by Qubit within 24 hours of DNA extraction. Presence of the 8.0 and 9.5 kb bands were assessed based on a single PCR reaction visualized on a 0.7% agarose gel. Stations are presented in the order in which they were sampled in the field. | | | | | | | |
| --- | --- | --- | --- | --- | --- | --- | --- |
| **Station** | **Latitude** | **Longitude** | **Biological Replicate** | **Volume Filtered (L)** | **Total DNA conc. (ng/μL)** | **9.5 kb band** | **8.0 kb band** |
| P28 | 47.7034 | -122.4544 | A | 2 | 3.75 | y | y |
|  |  |  | B | 2 | 3.78 | y | y |
|  |  |  | C | 2 | 3.4 | y | y |
| P5 | 47.8839 | -122.3675 | A | 2 | 3.64 | y | y |
|  |  |  | B | 2 | 3.63 | n | y |
|  |  |  | C | 2 | 4.29 | y | y |
| P1 | 48.0165 | -122.3042 | A | 2 | 3.71 | y | y |
|  |  |  | B | 2 | 6.21 | y | y |
|  |  |  | C | 2 | 4.94 | y | y |
| P3 | 48.1076 | -122.4896 | A | 2 | 4.21 | y | y |
|  |  |  | B | 2 | 4.91 | y | y |
|  |  |  | C | 2 | 4.68 | y | y |
| P4 | 48.2422 | -122.5533 | A | 2 | 5.31 | y | y |
|  |  |  | B | 2 | 5.02 | y | y |
|  |  |  | C | 2 | 5.11 | y | y |
| P26 | 48.3752 | -122.7167 | A | 2 | 5.26 | n | n |
|  |  |  | B | 2 | 7.74 | n | y |
|  |  |  | C | 2 | 6.81 | y | y |
| P22 | 48.2717 | -123.0189 | A | 2 | 3.14 | y | n |
|  |  |  | B | 2 | 2.63 | n | n |
|  |  |  | C | 2 | 3.82 | n | n |
| P21 | 48.1883 | -122.8504 | A | 2 | 7.05 | y | n |
|  |  |  | B | 2 | 9.31 | y | y |
|  |  |  | C | 2 | 13.4 | y | y |
| P20 | 48.142 | -122.6848 | A | 2 | 6.07 | y | y |
|  |  |  | B | 2 | 13 | n | y |
|  |  |  | C | 2 | 22.6 | y | y |
| P7 | 47.9835 | -122.6201 | A | 2 | 7.15 | y | y |
|  |  |  | B | 2 | 7.46 | y | y |
|  |  |  | C | 2 | 9.53 | y | y |
| P8 | 47.8967 | -122.6053 | A | 2 | 5.93 | y | n |
|  |  |  | B | 2 | 5.47 | n | n |
|  |  |  | C | 2 | 7.12 | y | n |
| P10 | 47.8001 | -122.7198 | A | 2 | 8.5 | n | y |
|  |  |  | B | 2 | 12.9 | y | y |
|  |  |  | C | 2 | 12.8 | y | y |
| P17 | 47.7356 | -122.7614 | A | 2 | 6.71 | y | y |
|  |  |  | B | 2 | 7.9 | y | y |
|  |  |  | C | 2 | 5.58 | n | n |
| P15 | 47.6616 | -122.8601 | A | 2 | 4.79 | n | n |
|  |  |  | B | 2 | 4.2 | n | n |
|  |  |  | C | 2 | 3.12 | y | n |
| P13 | 47.5471 | -123.008 | A | 2 | 2.94 | n | y |
|  |  |  | B | 2 | 5.5 | y | y |
|  |  |  | C | 2 | 4.67 | n | n |
| P12 | 47.4253 | -123.1083 | A | 2 | 7.47 | y | n |
|  |  |  | B | 2 | 4.16 | n | y |
|  |  |  | C | 2 | 4.97 | y | n |
| P402 | 47.3567 | -123.0233 | A | 2 | 14.2 | n | y |
|  |  |  | B | 2 | 14.2 | y | y |
|  |  |  | C | 2 | 9.48 | y | y |
| P29 | 47.5568 | -122.4433 | A | 1 | 1.64 | n | n |
|  |  |  | B | 1 | 1.81 | n | y |
|  |  |  | C | 1 | 2.05 | n | n |
| P30 | 47.4565 | -122.4084 | A | 2 | 3.46 | y | n |
|  |  |  | B | 2 | 3.6 | y | y |
|  |  |  | C | 2 | 3.34 | y | n |
| P31 | 47.3937 | -122.3601 | A | 1 | 1.78 | n | n |
|  |  |  | B | 1 | 1.56 | n | n |
|  |  |  | C | 1 | 1.83 | n | n |
| P33 | 47.3198 | -122.5008 | A | 2 | 3.82 | n | n |
|  |  |  | B | 2 | 4.21 | y | n |
|  |  |  | C | 2 | 4.84 | n | y |
| P35 | 47.1813 | -122.634 | A | 2 | 9.06 | n | y |
|  |  |  | B | 2 | 8.2 | n | y |
|  |  |  | C | 2 | 7.59 | n | n |
| P36 | 47.1684 | -122.7865 | A | 2 | 7.72 | y | n |
|  |  |  | B | 2 | 7.61 | n | y |
|  |  |  | C | 2 | 8.08 | n | y |
| P38 | 47.2766 | -122.7082 | A | 2 | 14.8 | n | y |
|  |  |  | B | 2 | 11.2 | y | y |
|  |  |  | C | 2 | 11.1 | y | y |

| Table S2. Species expected in the Seattle Aquarium 'Window on Washington Waters' tank, according to the Aquarium records. Species are categorized as fish (Teleostei) or non-fish (invertebrate); Tank census: number of expected individuals in the tank for vertebrates (not recorded for non-vertebrate species); eDNA detection: binary, observed in LR-PCR from eDNA or not. eDNA detection is not reported for non-fish species, as our annotation database only included vertebrates. | | | | |
| --- | --- | --- | --- | --- |
| **Fish (Y/N)** | **Common name** | **Scientific name** | **Tank census (#)** | **eDNA detection** |
| Y | Pacific herring | *Clupea pallasii* | 0 | Y |
| Y | Pink salmon | *Oncorhynchus gorbuscha* | 0 | Y |
| Y | Redbanded rockfish | *Sebastes babcocki* | 0 | Y |
| Y | English sole | *Parophrys vetulus* | 0 | Y |
| Y | C-O sole | *Pleuronichthys coenosus* | 0 | Y |
| Y | Canary rockfish | *Sebastes pinniger* | 41 | Y |
| Y | Coho salmon | *Oncorhynchus kisutch* | 60 | Y |
| Y | Copper rockfish | *Sebastes caurinus* | 11 | Y |
| Y | Kelp greenling | *Hexagrammos decagrammus* | 5 | N |
| Y | Red Irish lord | *Hemilepidotus hemilepidotus* | 5 | Y |
| Y | Wolfeel | *Anarrhichthys ocellatus* | 5 | Y |
| Y | Yelloweye rockfish | *Sebastes ruberrimus* | 4 | Y |
| Y | Yellowtail rockfish | *Sebastes flavidus* | 21 | Y |
| Y | Bocaccio rockfish | *Sebastes paucispinis* | 2 | N |
| Y | Black rockfish | *Sebastes melanops* | 13 | Y |
| Y | Black-eyed goby | *Rhinogobiops nicholsii* | 1 | N |
| Y | Brown Irish lord | *Hemilepidotus spinosus* | 1 | N |
| Y | Cabezon | *Scorpaenichthys marmoratus* | 1 | N |
| Y | China rockfish | *Sebastes nebulosus* | 8 | N |
| Y | Deacon rockfish | *Sebastes mystinus* | 24 | N |
| Y | Quillback rockfish | *Sebastes maliger* | 7 | N |
| Y | Rock greenling | *Hexagrammos lagocephalus* | 1 | N |
| Y | Rosy rockfish | *Sebastes rosaceus* | 1 | N |
| Y | Tiger rockfish | *Sebastes nigrocinctus* | 5 | N |
| Y | Widow rockfish | *Sebastes entomelas* | 4 | N |
| N | California sea cucumber |  | nr | nr |
| N | Fish-eating anemone |  | nr | nr |
| N | Green anemone |  | nr | nr |
| N | Green sea urchin |  | nr | nr |
| N | Gumboot chiton |  | nr | nr |
| N | Kelp crab |  | nr | nr |
| N | Keyhole limpet |  | nr | nr |
| N | Leather star |  | nr | nr |
| N | Mossy chiton |  | nr | nr |
| N | Pacific blood star |  | nr | nr |
| N | Painted anemone |  | nr | nr |
| N | Plate limpet |  | nr | nr |
| N | Plumose anemone |  | nr | nr |
| N | Purple-ringed top snail |  | nr | nr |
| N | Rainbow star |  | nr | nr |
| N | Red Sea urchin |  | nr | nr |
| N | Velcro star |  | nr | nr |
| N | Vermillion star |  | nr | nr |
| N | White-spotted rose anemone |  | nr | nr |
| N | Wrinkled dogwinkle |  | nr | nr |

**SUPPLEMENTAL FIGURES**

**Figure S1. Gel electrophoresis images showing the presence or absence of target fragments after LR-PCR.** The 9.5 kb (**A, B**) and 8.0 kb (**C, D**) target bands obtained after LR-PCR amplification of environmental samples collected during the April 2024 Salish Sea cruise. (**E)** The 9.5 and 8.0 kb target bands obtained after LR-PCR of mesocosm samples collected in August of 2024 from the Seattle Aquarium. Prior to loading, 3 μL of each post-PCR reaction was mixed with 2 μL loading dye containing 1X SYBR Green. Each gel was comprised of 0.7% agarose and run at 3-4 volts/cm for approximately 120 minutes. 2 μL undiluted Quick-Load 1 kb Extend DNA Ladder (New England Biolabs) mixed with 2 μL loading dye containing 1X SYBR Green was used as a reference.

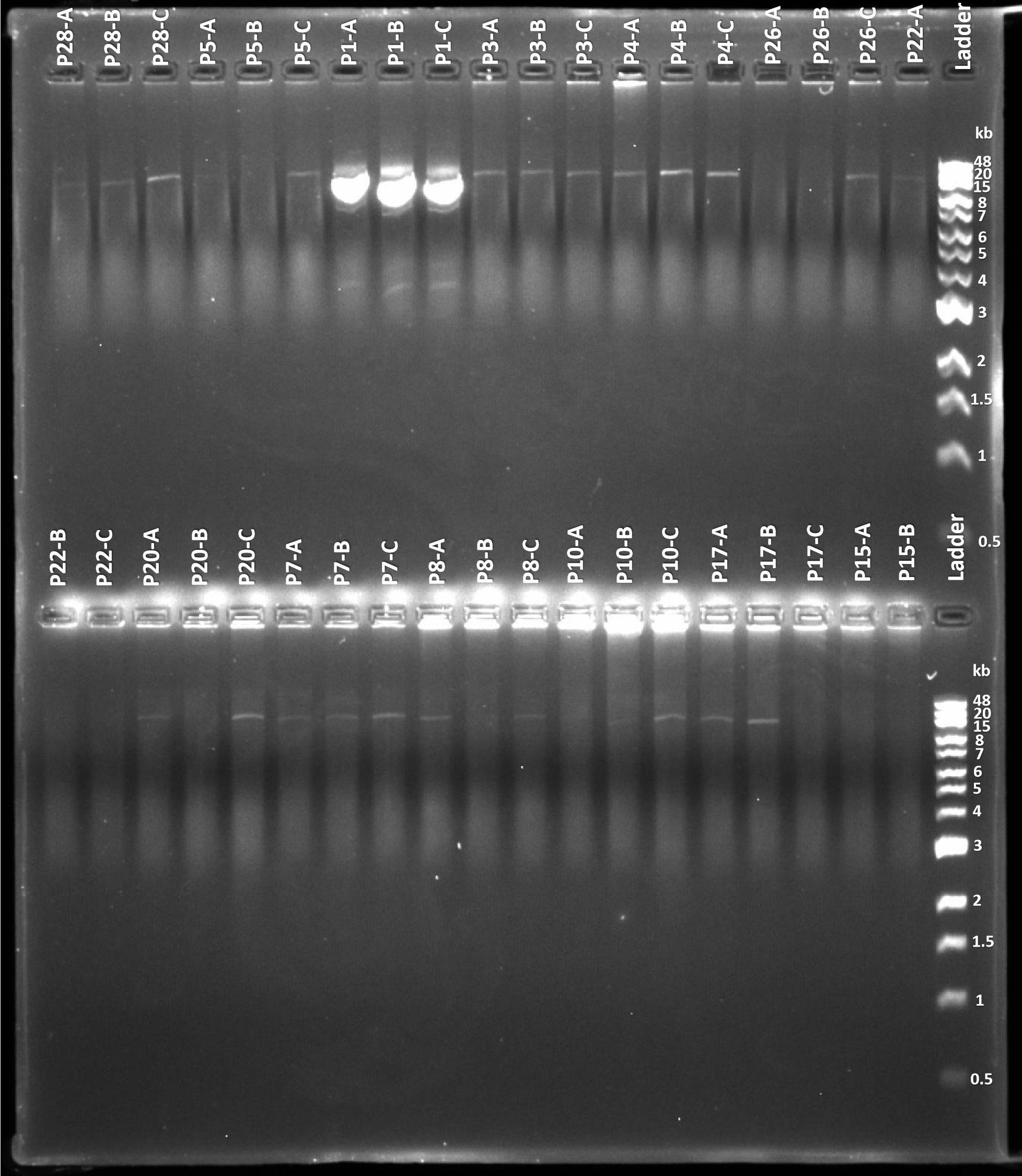
A.

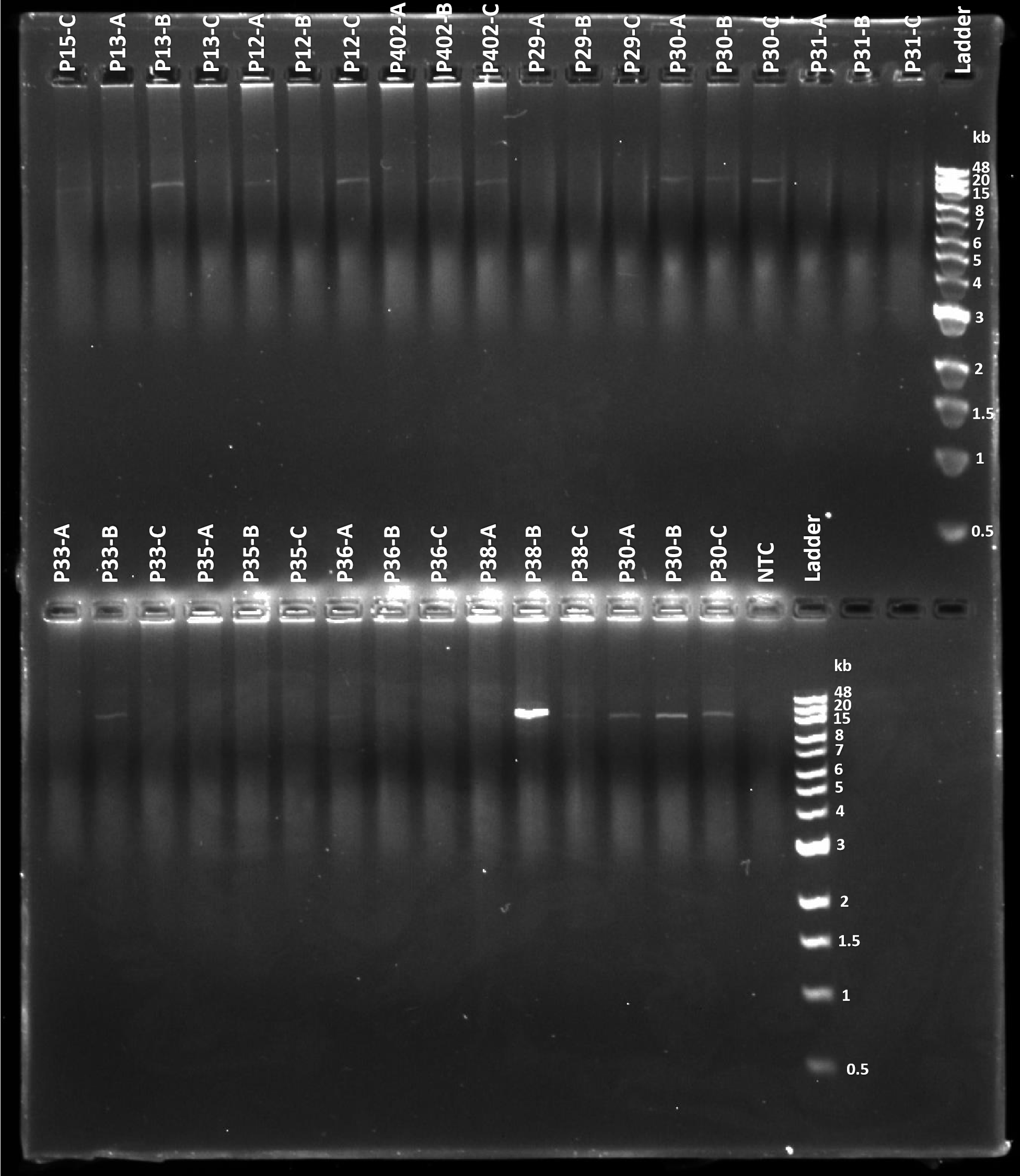
B.

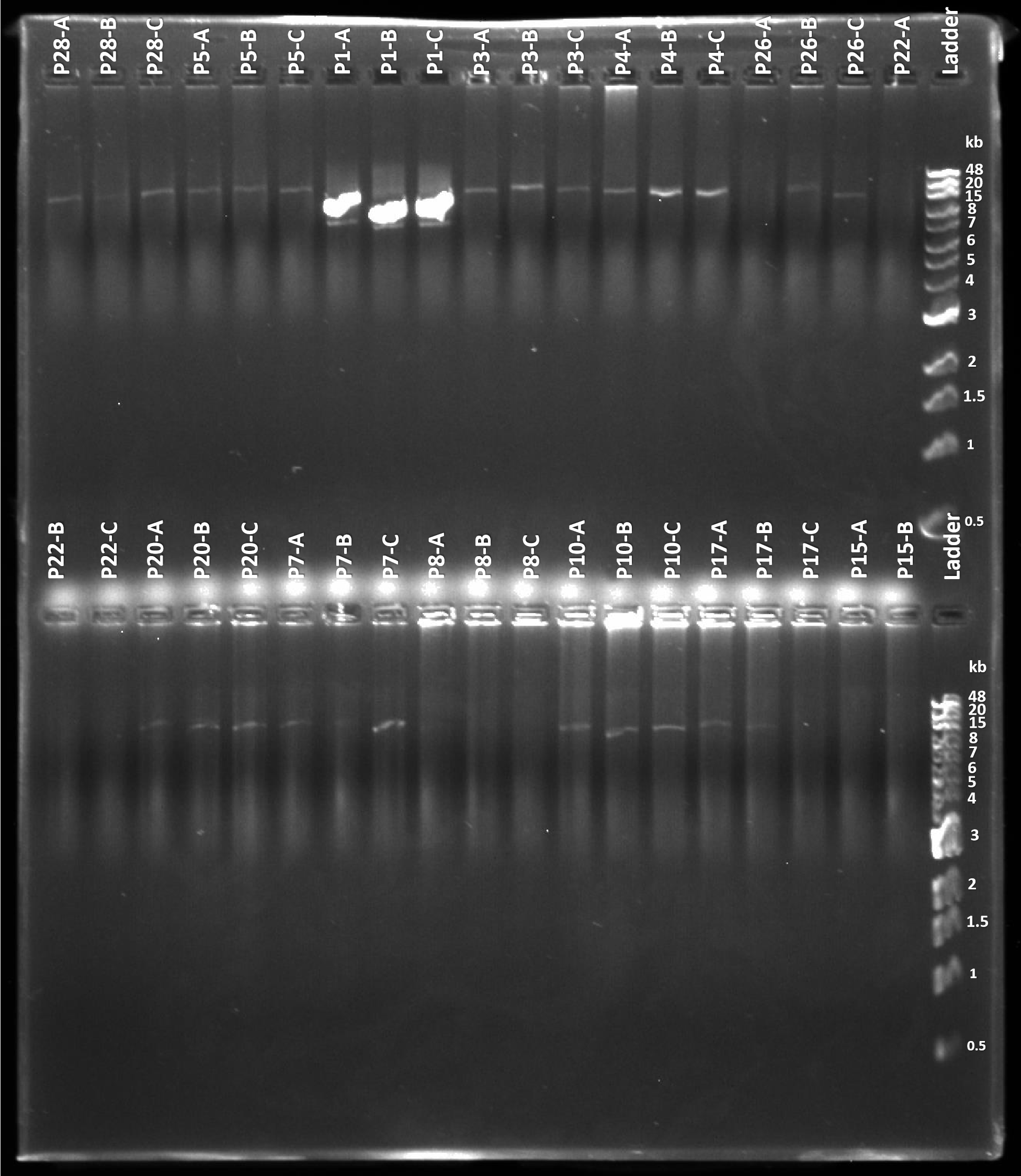
C.

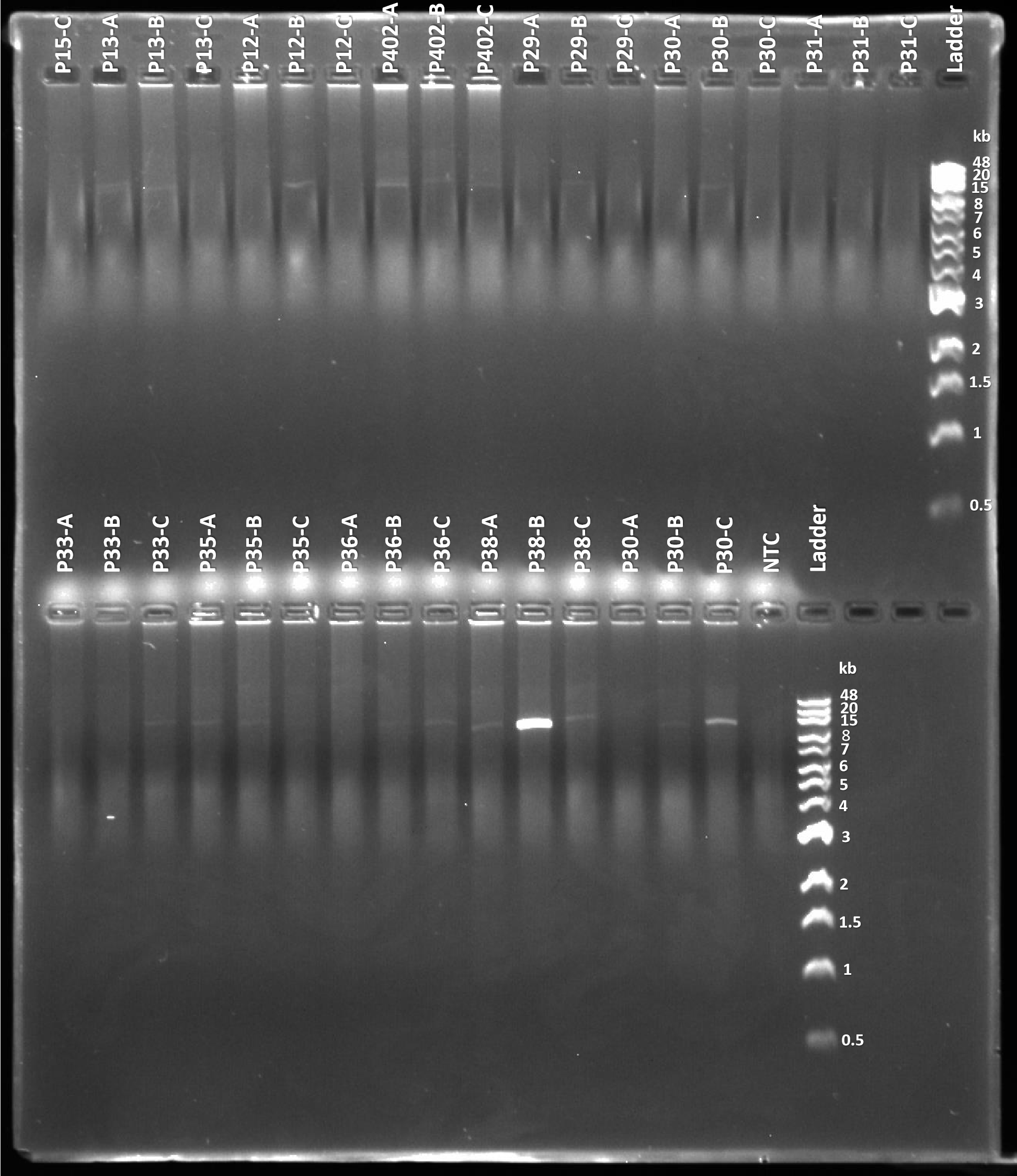
D.

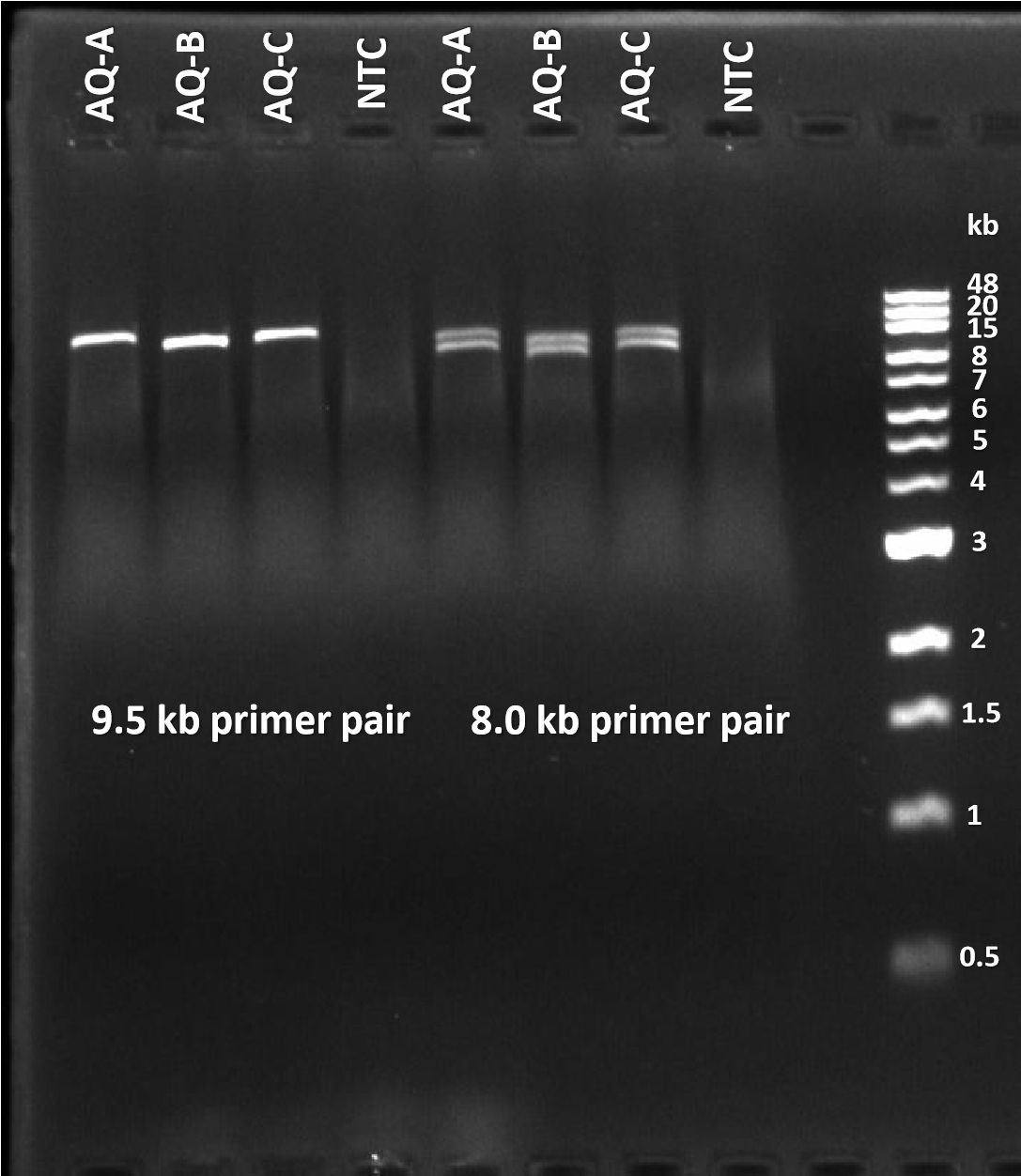
E.

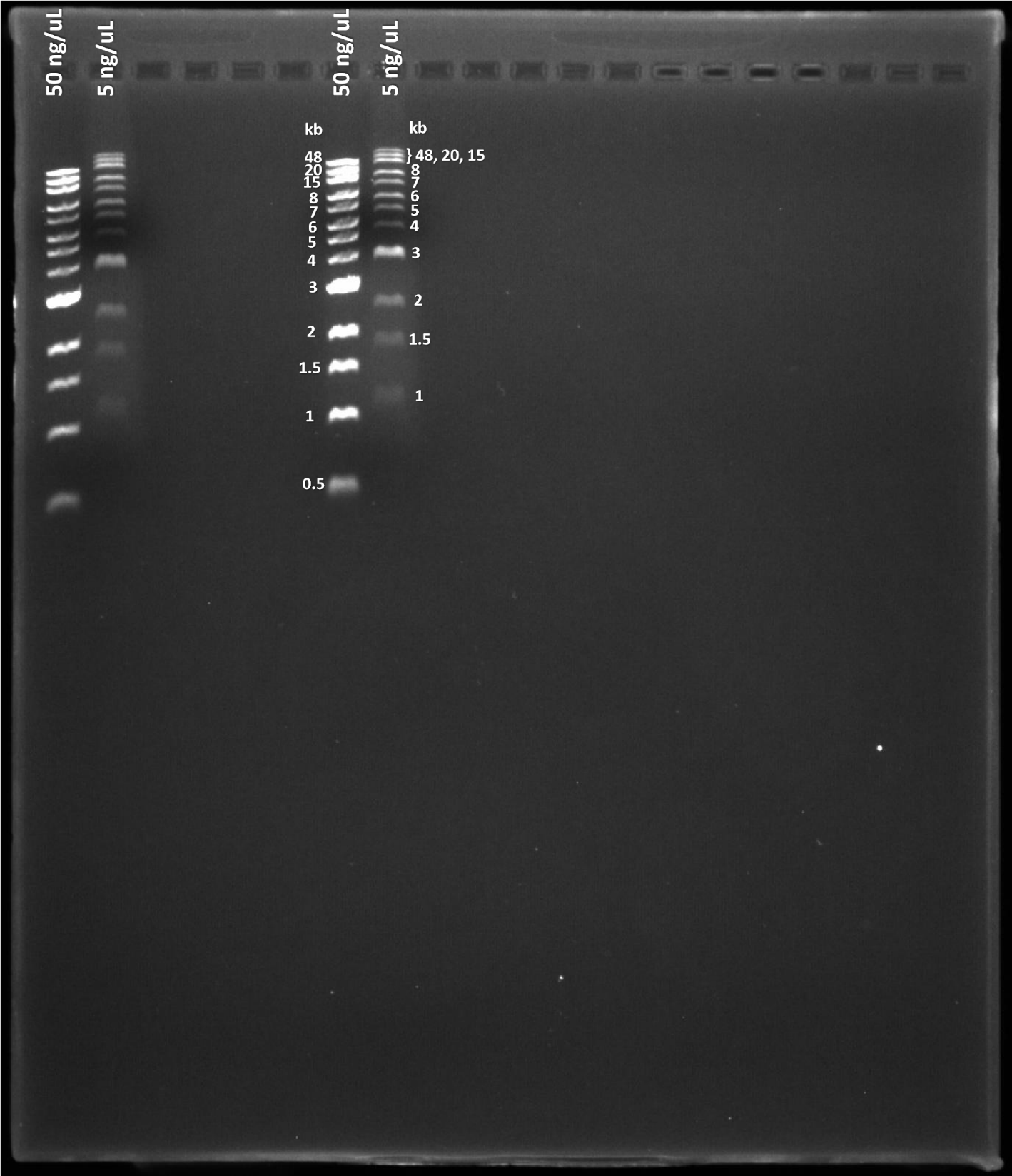
**Figure S2. Gel electrophoresis image showing differential migration of reference ladder, dependent on the loading concentration.** Prior to loading, 2 μL of either stock concentration (50 ng/μL) or a 1:10 dilution (5 ng/μL) of Quick-Load 1 kb Extend DNA Ladder (New England Biolabs), was mixed with 2 μL loading dye containing 1X SYBR Green. The gel contained 0.7% agarose and was run at 3-4 volts/cm for 120 minutes.
